# Recombining your way out of trouble: The genetic architecture of hybrid fitness under environmental stress

**DOI:** 10.1101/622845

**Authors:** Zebin Zhang, Devin P. Bendixsen, Thijs Janzen, Arne W. Nolte, Duncan Greig, Rike Stelkens

**Author notes:** These authors contributed equally to the study (Zebin Zhang, Devin P. Bendixsen). Corresponding author: Rike Stelkens, Department of Zoology, Stockholm University, Svante Arrheniusväg 18 B, 106 91 Stockholm, Sweden.

## Abstract

Hybridization between species is a fundamental evolutionary force that can both promote and delay adaptation. There is a deficit in our understanding of the genetic basis of hybrid fitness, especially in non-domesticated organisms. We also know little about how hybrid fitness changes as a function of environmental stress. Here, we made genetically variable F2 hybrid populations from two divergent *Saccharomyces* yeast species, exposed populations to ten toxins, and sequenced the most resilient hybrids on low coverage using ddRADseq. We expected to find strong negative epistasis and heterozygote advantage in the hybrid genomes. We investigated three aspects of hybridness: 1) hybridity, 2) interspecific heterozygosity, and 3) epistasis (positive or negative associations between non-homologous chromosomes). Linear mixed effect models revealed strong genotype-by-environment interactions with many chromosomes and chromosomal interactions showing species-biased content depending on the environment. Against our predictions, we found extensive selection against heterozygosity such that homozygous allelic combinations from the same species were strongly overrepresented in an otherwise hybrid genomic background. We also observed multiple cases of positive epistasis between chromosomes from opposite species, confirmed by epistasis- and selection-free simulations, which is surprising given the large divergence of the parental species (~15% genome-wide). Together, these results suggest that stress-resilient hybrid genomes can be assembled from the best features of both parents, without paying high costs of negative epistasis across large evolutionary distances. Our findings illustrate the importance of measuring genetic trait architecture in an environmental context when determining the evolutionary potential of hybrid populations.

## Introduction

Populations exposed to gene flow, introgression or hybridization contain vast amounts of genetic variation, sometimes producing phenotypes with more extreme adaptations than found in the parent populations (Shahid, et al. 2008; Stelkens and Seehausen 2009; Pritchard, et al. 2013; Stelkens, Brockhurst, Hurst, Miller, et al. 2014; Holzman and Hulsey 2017). The average fitness of hybrid crosses is usually lower than that of non-hybrid crosses due to interspecific incompatibilities or other negative effects, e.g. the break-up and loss of co-adapted beneficial gene complexes for local adaptation (Coyne and Orr 2004). This applies especially to the hybrid offspring of genetically divergent lineages. However, some hybrids show high fitness (Rieseberg, et al. 1999; Dittrich-Reed and Fitzpatrick 2012), which is exploited in agricultural breeding to improve the yield, taste or other desirable traits of cultivars (e.g. Marullo, et al. 2006; Kuczyńska, et al. 2007; Shivaprasad, et al. 2012; Koide, et al. 2019). High hybrid fitness can also be relevant for macroevolutionary dynamics and the generation of biodiversity, when ecologically divergent hybrid phenotypes form their own evolutionary lineage and become reproductively isolated from the parents (Anderson and Stebbins 1954; Lewontin and Birch 1966; Rieseberg, et al. 2003; Seehausen 2004; Arnold 2006; Mallet 2007; Nolte and Tautz 2010; Abbott, et al. 2013; Schumer, et al. 2014).

Experimental evolution with interspecific hybrids of the budding yeast *Sacchromyces spp*. has demonstrated that hybrids can colonize new ecological niches (Stelkens, Brockhurst, Hurst and Greig 2014; Stelkens, Brockhurst, Hurst, Miller, et al. 2014), and gain resistance to stress (Greig, et al. 2002; Piotrowski, et al. 2012; Dunn, et al. 2013). However, the genetic mechanisms allowing some hybrids to express high fitness phenotypes in the face of negative epistasis between divergent parental genomes are largely unknown, especially in non-domesticated organisms. Also, we know very little about the impact of stressful and deteriorating environmental conditions on the evolutionary potential of hybrid populations, although this is becoming increasingly relevant in the face of global environmental change. So far empirical studies of hybrid fitness in an environmental context are scant and the results are mixed. Some found environmental stress to intensify the negative effects of hybridization (Koevoets, et al. 2012; Barreto and Burton 2013), others found hybrid fitness to increase with stress (Edmands and Deimler 2004; Willett 2012; Hwang, et al. 2016b); again others detected no effects of stress on hybrid fitness (Armbruster, et al. 1999).

Here, we made genetically highly variable F2 hybrids by crossing two divergent species of *Saccharomyces* (*S. cerevisiae* and *S. paradoxus*). These species have well-sequenced, well-assembled genomes that differ at ~15% of nucleotides genome-wide (Liti, et al. 2009). Due to their large divergence these species produce only ~1% viable F2 hybrid offspring (Hunter, et al. 1996; Boynton, et al. 2018; Rogers, et al. 2018). Because of this, we expected to find strong negative epistasis between genetic material from opposite species and, potentially, heterozygote advantage in the genomes of viable hybrids. We exposed 240 populations of these viable F2 hybrids to ten stressful environments containing high concentrations of different toxins (e.g. caffeine, ethanol, lithium acetate; Table S1). At the end of the growth period, we sequenced the genomes of 240 hybrid survivors at low coverage using double digest, restriction-site associated DNA (ddRAD) markers and mapped hybrid genomes to both reference genomes. To identify the genetic and environmental factors shaping hybrid genomes in our experiment, we measured three aspects of ‘hybridness’; 1) hybridity (proportion of hybrid genome mapping to one or other parent species); 2) interspecific heterozygosity (homologous chromosomes from opposite species), and 3) epistasis (positive or negative associations between non-homologous chromosomes from opposite species).

Hybridity is a continuous measure, e.g. the proportion of genomic material inherited from one over the other parental species (Gompert and Buerkle 2016), often referred to as hybrid index (Barton and Gale 1993; Buerkle 2005) or admixture proportion (Pritchard, et al. 2000; Falush, et al. 2003). Hybridity is useful to place hybrid individuals on a single axis ranging from zero to one with pure parental individuals at opposite ends. However, this simple measure can mask potentially important fitness effects of individual loci (or chromosomes) deviating from the average hybridity of the genome. For instance, dominance (the reciprocal masking of deleterious alleles at multiple loci; Bruce 1910)) and overdominance (the intrinsic benefit of being heterozygous for at least one locus) can produce hybrids with high fitness. This is known as ‘interspecific heterozygosity’ or ‘inter-population ancestry’ (Barton 2000; Fitzpatrick 2012; Lindtke, et al. 2012; Gompert and Buerkle 2013). As an example, while every chromosome in a diploid F1 hybrid is heterozygous, maximizing within-locus hybridity, the genome still carries a full haploid set of both parental chromosomes. At the same time, a diploid F2 or higher generation hybrid can, theoretically, be composed of fully homozygous chromosomes (i.e. homologous chromosomes are from the same species), which minimizes within-locus hybridity. But this F2 hybrid may contain half the chromosomal set from either parent, maximizing between-locus hybridity. Because these types of hybrids would be indistinguishable with a single hybrid index (which would be 0.5 in both examples), we also measured interspecific heterozygosity, i.e. chromosomes in the F2 hybrid genome carrying interspecific, homologous combinations.

Epistasis is caused by interactions between alleles from at least two different loci increasing or decreasing fitness more than the sum of the individual contributions of these loci. Negative epistasis is the core element of the Dobzhansky-Muller model of genetic incompatibilities often recruited to explain the evolution of reproductive isolation and outbreeding depression between biological species, with increasing negative impact the more divergent the parental genomes are (Dobzhansky 1936; Müller 1942; Lynch 1991). Given the large nucleotide divergence between the parental lineages used here (~15% genome-wide), we expected negative epistasis to be prominent in F2 hybrids, determining variation in fitness to a large extent (Cubillos, et al. 2011; Shapira, et al. 2014). We measured epistasis by testing for significant associations (presence and absence) between non-homologous chromosomes from different species in F2 hybrid yeast genomes, and compared our experimental data to computer-simulated data, assuming free segregation of chromosomes, no epistasis and no selection in a theoretical F2 hybrid population.

We also predicted species biases on individual chromosomes and associations between chromosomes, and predicted species biases to be determined by environmental changes (Kvitek and Sherlock 2011; de Vos, et al. 2013). For instance, Jaffe et al. (2019) showed that adding more environmental conditions tripled the number of genetic interactions detected in fitness assays between double mutants of yeast (Jaffe, et al. 2019). Filteau et al. (2015) found that the course of compensatory evolution rescuing yeast populations from the negative fitness effects of deleterious mutations was strongly contingent on environmental context (different carbon sources) (Filteau, et al. 2015). Similarly, Lee et al., dissecting the genetic basis of a yeast colony phenotype, found that environmental stress affected the impact of epistasis, additivity, and genotype-environment interactions (Lee, et al. 2019). We tested for species biases and genotype-by-environment interactions using linear mixed effect models.

## Results

We sequenced a total of 240 F2 hybrid strains, of which 184 were mostly diploid and 53 were mostly haploid (three genomes were discarded due to low read quality). Thus, the majority of spores germinated and mated in the 96-well plates, forming diploid F2 hybrids homozygous for both drug resistance markers *cyh2r* and *can1r*. The haploid genotypes detected by sequencing were unmated F2 (i.e. F1 spores), and hemizygous for *cyh2r* and *can1r*. Occasional chromosomal copy number variation within genomes was apparent, but variation in read depth along chromosomes did not allow for a systematic quantification of disomies (or higher - somies). Because we saw no significant differences between F2 samples isolated from high concentration of a toxin and low concentration in any of the tests, we proceeded by pooling all samples for analysis.

### Interspecific heterozygosity

Assuming no selection and free segregation of chromosomes, interspecific heterozygosity (i.e. having homologous chromosomes from both species) was lower than expected in the 187 diploid F2 genomes across all toxins (mean heterozygosity = 0.44 median = 0.36; Figure 1A). On average, only 7 of the 16 chromosomes per genome carried a heterospecific combination. In total, there were almost 1.3 times more homozygous chromosomes (n = 1450; 57%) than heterozygous chromosomes (n = 1126; 43%). Environments differed substantially in the proportions of homo- and heterozygous chromosomes (Figure 1B). While in some environments, F2 hybrids were mainly homozygous (e.g. zinc sulfate), other environments promoted the growth of mainly heterozygote genotypes (e.g. NaCl). This was confirmed using simulations. Five stress environments (zinc sulfate, citric acid, ethanol, salicylic acid and caffeine) produced mean interspecific heterozygosities below the range expected based on random segregation simulations (Figure S1). Seven environments produced mean homozygosities above the expected range: Three environments (zinc sulfate, citric acid and ethanol) produced F2 genomes more homozygous than the simulated range of expected homozygosity for *S. cerevisiae* predicted. Four environments (caffeine, citric acid, salicylic acid and zinc sulfate) produced F2 genomes with higher homozygosity than the simulated range of expected homozygosity for *S. paradoxus*.

Testing for variation in zygosity between chromosomes, across all environments, we found three chromosomes with a significantly larger proportion of *S. cerevisiae* homozygotes (chromosomes 4, 5, 14), and three chromosomes with a significantly larger proportion of *S. paradoxus* homozygotes (chromosomes 6, 7, 10; Figure 1C). Note that chromosomes 5 and 7 carried resistance genes, which, as predicted, selected them to be homozygous for *S. cerevisiae* and *S. paradoxus*, respectively (chromosome 5 also carried some *S. paradoxus* content due to recombination at one end of the chromosome). Three chromosomes were more often heterozygous than expected by chance (chromosomes 2, 3, 12).

Using a linear mixed-effect model to understand what causes variation in zygosity between environments, chromosomes and genomes, we found that the interaction between *chromosome ID* and *environment* best predicted zygosity (Table S2). Thus, which chromosome was homozygous, and which chromosome was heterozygous, was largely determined by the stress environment from which it was sampled.

**Figure 1.**
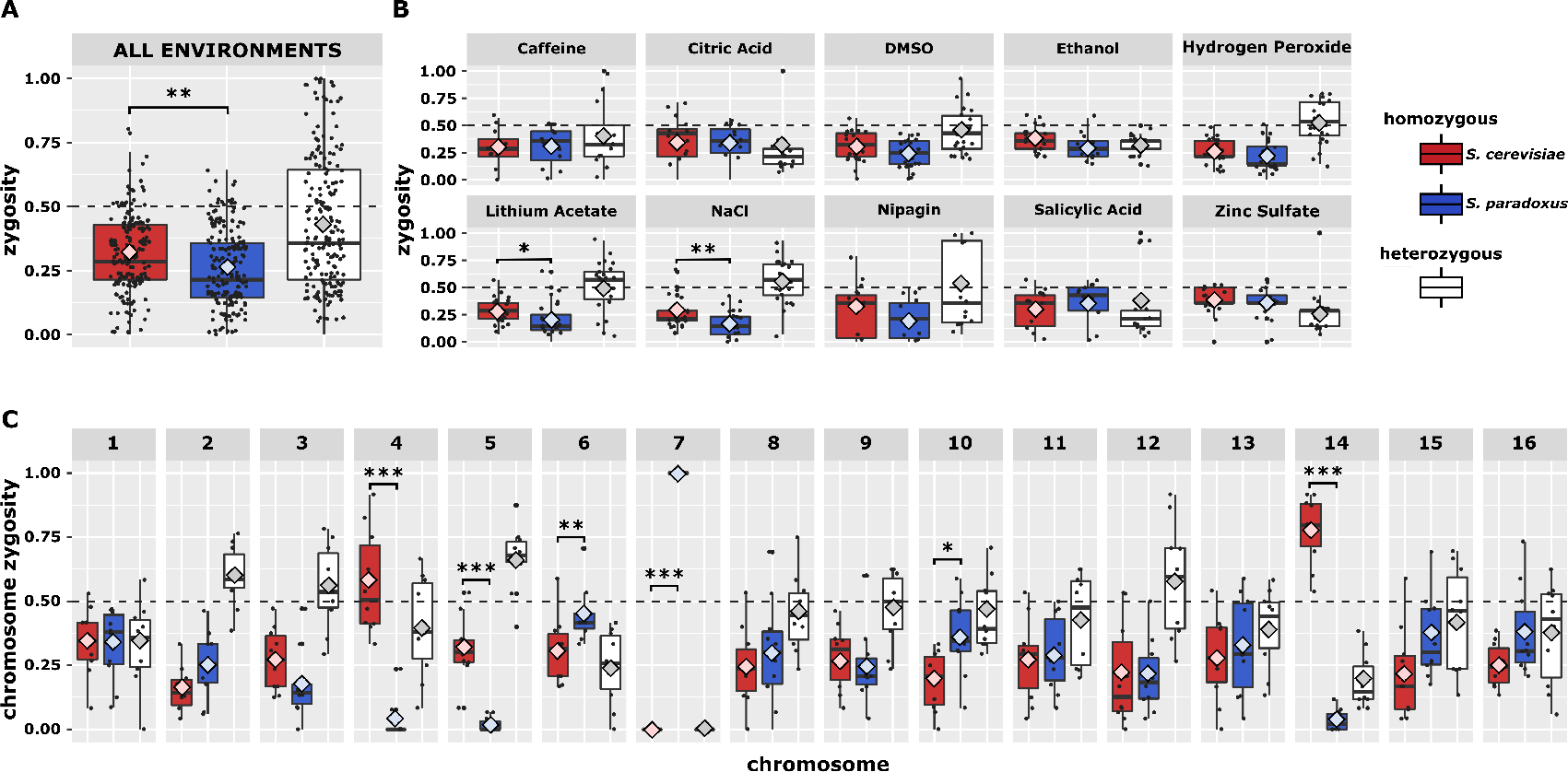
Homo-/heterozygosity of diploid F2 hybrid genomes. **(A)** Mean zygosity of 187 diploid F2 hybrids across all chromosomes (except chromosome 5 and 7) and across all environments. Dashed horizontal line (at 0.5) shows expected heterozygosity without selection and free segregation. Black lines in boxes are medians and large diamonds indicate means. Each black dot represents a genome. Boxes are interquartile ranges (IQR). Whiskers are 1.5 x IQR **(B)** Mean F2 hybrid zygosity per environment. **(C)** Mean F2 hybrid zygosity per chromosome. Chromosome 5 and 7 harbour drug markers selecting for *S. cerevisiae* and *S. paradoxus* chromosomes, respectively. Each dot represents one of 10 environments. Asterisks indicate significant differences in proportions of *S. cerevisiae* vs. *S. paradoxus* homozygotes in Wilcoxon signed-rank tests (p = *0.05, **0.01, ***0.001).

### Genome-wide hybridity

Average genome-wide hybridity (defined to be % markers inherited from *S. cerevisiae*) was 0.52 ± 0.05 (μ±SD) across all chromosomes and all environments (Figure S2A). Thus, genomic contributions were roughly balanced between the two parental species with a slight overrepresentation of *S. cerevisiae*. The most biased genome towards *S. cerevisiae* was 0.68 and the most biased towards *S. paradoxus* was 0.35. Average hybridity per toxin ranged from 0.49 ± 0.07 in salicylic acid (*paradoxus*-biased) to 0.55 ± 0.07 (*cerevisiae*-biased) in Nipagin, but no significant differences were found between toxins (Figure S2B). The distributions and means of genome-wide hybridity observed in each environment were all within the range expected from simulations based on random chromosome segregation (Figure S3). Chr. 5 and 7 were excluded from these analyses because they carried antibiotic resistance markers selecting for *S. cerevisiae and S. paradoxus* hemi- or homozygosity, respectively.

### Chromosome hybridity

A closer examination of hybridity (defined to be % markers inherited from *S. cerevisiae*) at the chromosomal level revealed species biases in inheritance patterns (Figure 2A). For most chromosomes the average chromosome hybridity across all environments was approximately 0.5, suggesting equal inheritance from *S. cerevisiae* and *S. paradoxus*. As expected chromosome 5 was inherited primarily from *S. cerevisiae* and chromosome 7 was inherited primarily from *S. paradoxus* because they carried species-specific drug markers. Interestingly, chromosome 4 and 14 were also inherited predominantly from *S. cerevisiae* (0.77 and 0.88 respectively). Nine of the 16 chromosomes exhibited varying levels of hybridity depending on environment (ANOVA, p < 0.05), with six showing significant variation (Figure 2B, p < 0.001). For some chromosomes, the hybridity variation resulting from environment did not change the species bias (chromosomes 7, 14). However, some chromosomes shifted from unbiased (~ 0.5 hybridity) to biased for *S. paradoxus* in some environments (chromosomes 10, 15). Two chromosomes exhibited environment-dependent species bias (chromosomes 12, 13). In some environments these chromosomes were biased towards *S. cerevisiae* (> 0.65%) and in other environments biased towards *S. paradoxus* (< 0.35%). Using a linear mixed-effect model we found that the interaction between *chromosome ID* and *environment* and the interaction between *chromosome ID* and *ploidy* (haploid or diploid) best predicted the hybridity of the chromosome (Table S3). The optimal model did not change if data from chromosome 5 and 7 were excluded.

**Figure 2.**
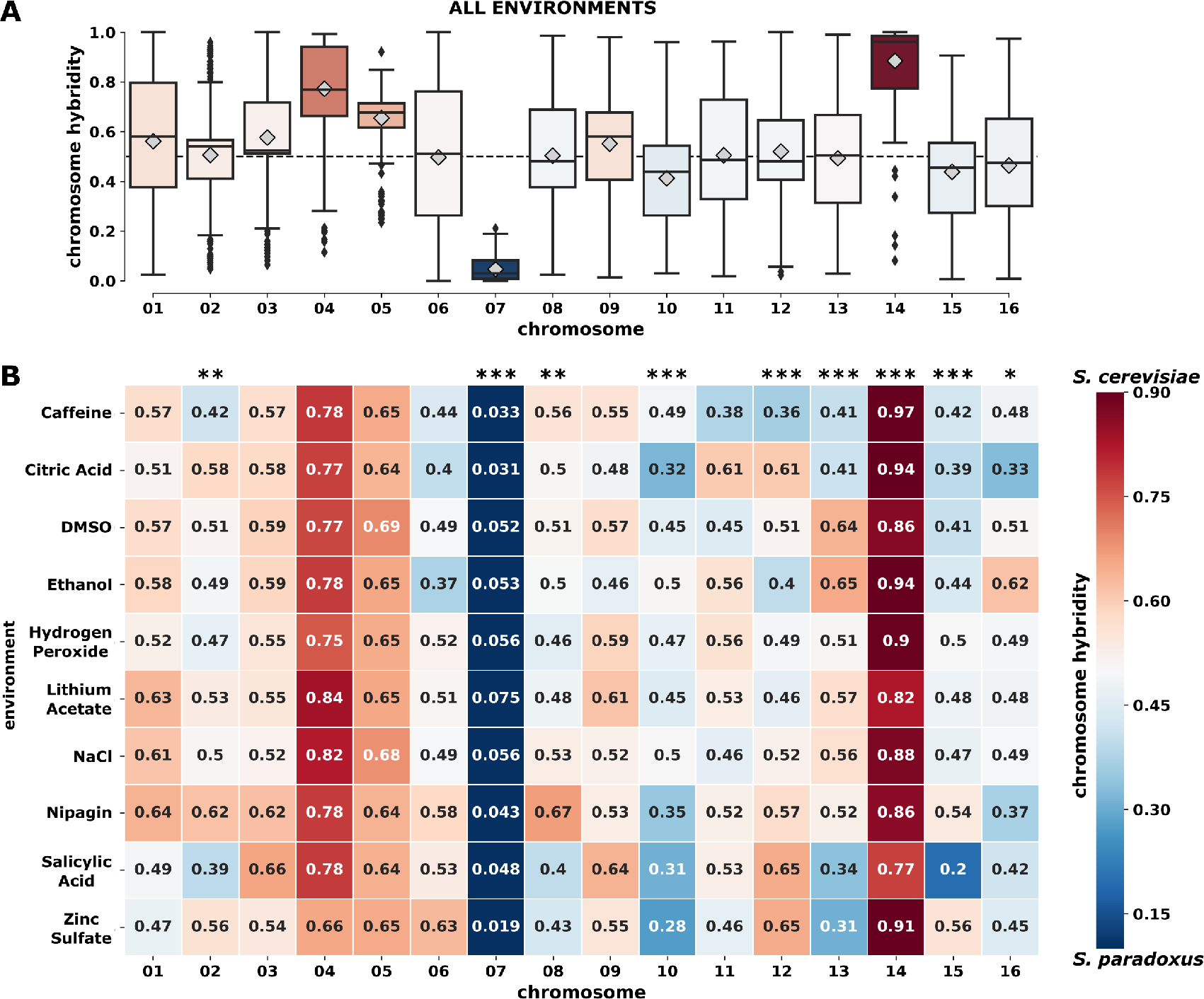
Chromosome hybridity of 237 haploid and diploid F2 genomes. (**A)** Chromosome hybridity for each chromosome across all environments. Chromosome hybridity is measured as the percent chromosome mapping to *S. cerevisiae*. Boxplots as in Figure 1, but colored according to where the median falls. Dashed line (0.5) indicates equal amounts of the chromosome mapping to *S. cerevisiae*. Species biases on chromosome 5 (~65% from *S. cerevisiae)* and 7 (~5% from *S. cerevisiae)* are by design and result from alternative recessive drug markers. **(B)** Chromosome hybridity for each chromosome within environments. Numbers in coloured boxes indicate average chromosome hybridity across 24 F2 diploid genomes. Asterisks indicate significant differences in average chromosome hybridity across environments (p = *0.05, **0.01, ***0.001).

### Chromosome hybridity interactions

Testing for chromosomal hybridity interactions within a given F2 genome revealed strong environment-dependent effects. A large difference in chromosome hybridity for each chromosome combination, defined as delta chromosome hybridity (DCH), indicated that two chromosomes mapped primarily to opposing species within a given F2 genome. This could potentially result from negative epistatic interactions within species (i.e. chromosome incompatibility), from positive epistatic interactions between species or a combination of the two. Some chromosomes maintained high DCH suggesting that they were preferably from opposing species across all environments (Figure 3A, chr7 x chr4, chr7 x chr14, chr7 x chr5). Chromosome 7 and 5 were designed to come from opposing species and were therefore expected to have high DCH. However, the remaining high DCH interactions with chromosome 7 suggest potential chromosome incompatibilities or strong positive epistatic interactions between species. Chromosome 7 was designed to be inherited from *S. paradoxus* and potentially resulting from this, chromosome 4 and chromosome 14 were primarily inherited from *S. cerevisiae*. Analyzing the DCH in each stress environment independently showed that different environments exhibited varying levels and distributions of DCH (Figure 3B). Two environments (zinc sulfate, salicylic acid) exhibited higher mean DCH than the range expected from simulations based on no chromosome interactions (Figure 4). Four environments (NaCl, hydrogen peroxide, lithium acetate, DMSO) exhibited lower mean DCH than the simulated expected range.

Analyzing directional correlations between hybridities of chromosomes within a genome revealed similar environment-dependent interactions (Figure 5). Significant positive correlations suggest positive epistatic interactions within species, meaning that as chromosome hybridity in one chromosome shifted towards one species (1 = *S. cerevisiae*, 0 = *S. paradoxus*), the chromosome hybridity in the linked chromosome also shifted towards the same species (Fig. 5A). Inversely, negative correlations suggested negative epistatic interactions within species. A lack of correlation suggested no significant epistatic interaction. For all environments combined and individually, the distribution of correlations was approximately normal with a mean near zero. There was a range of significant correlations (p < 0.01) found across environments ranging in numbers from 0 in salicylic acid and citric acid to 11 in NaCl (Figure 5B). The distributions and means for each environment all fell within the range that was expected from simulations based on no chromosome interactions (Figure S4). However, although the distributions and means were within the range simulated, three environments (caffeine, lithium acetate and NaCl) exhibited more significant correlations (p < 0.01) than expected (Figure S5). The NaCl environment resulted in a total of 11 significant hybridity correlations (8 positive, 3 negative), whereas 10,000 simulations never encountered more than 7 correlations by chance.

**Figure 3.**
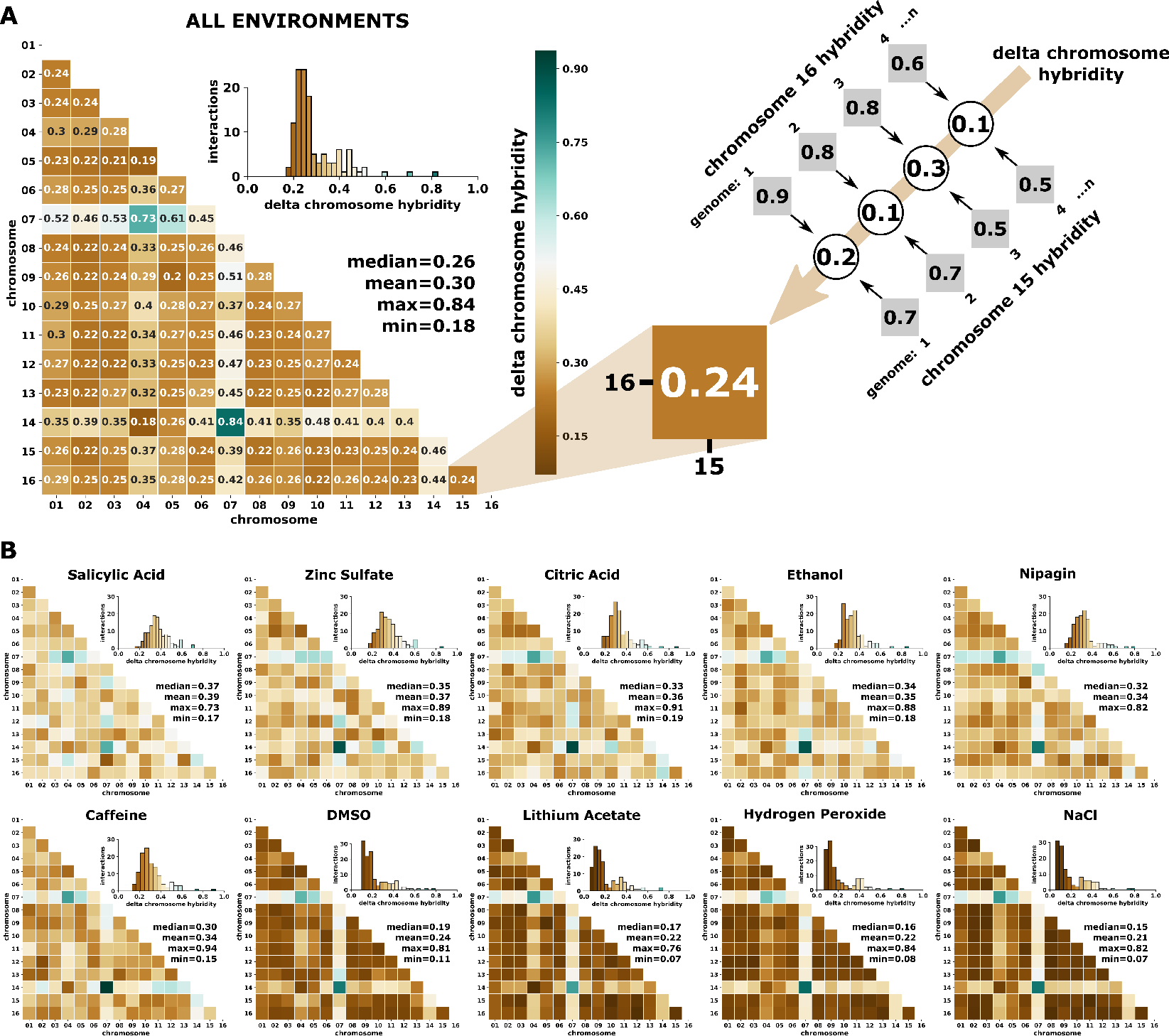
Interactions of chromosome hybridities altered by environment. **(A)** Average change in chromosome hybridity (percent chromosome mapping to *S. cerevisiae*) between chromosomes across all environments. Inset depicts the distribution of delta chromosome hybridity for all chromosome interactions. Delta chromosome hybridity is determined for each chromosome interaction by taking the difference between the hybridity measurements for each chromosome within an F2 genome (n genomes = 237). These values are then averaged across all diploid F2 genomes, and medians and means are reported and colored accordingly. A large delta chromosome hybridity (green) suggests that the chromosomes map primarily to opposing species. A small delta chromosome hybridity (brown) suggests that the chromosomes have similar levels of hybridity. These chromosomes may come from primarily the same species or at least have similar hybridity proportions. **(B)** Average change in chromosome hybridity between chromosomes for each environment. Calculations are performed the same as panel A for each environment independently (n genomes ~ 24).

**Figure 4.**
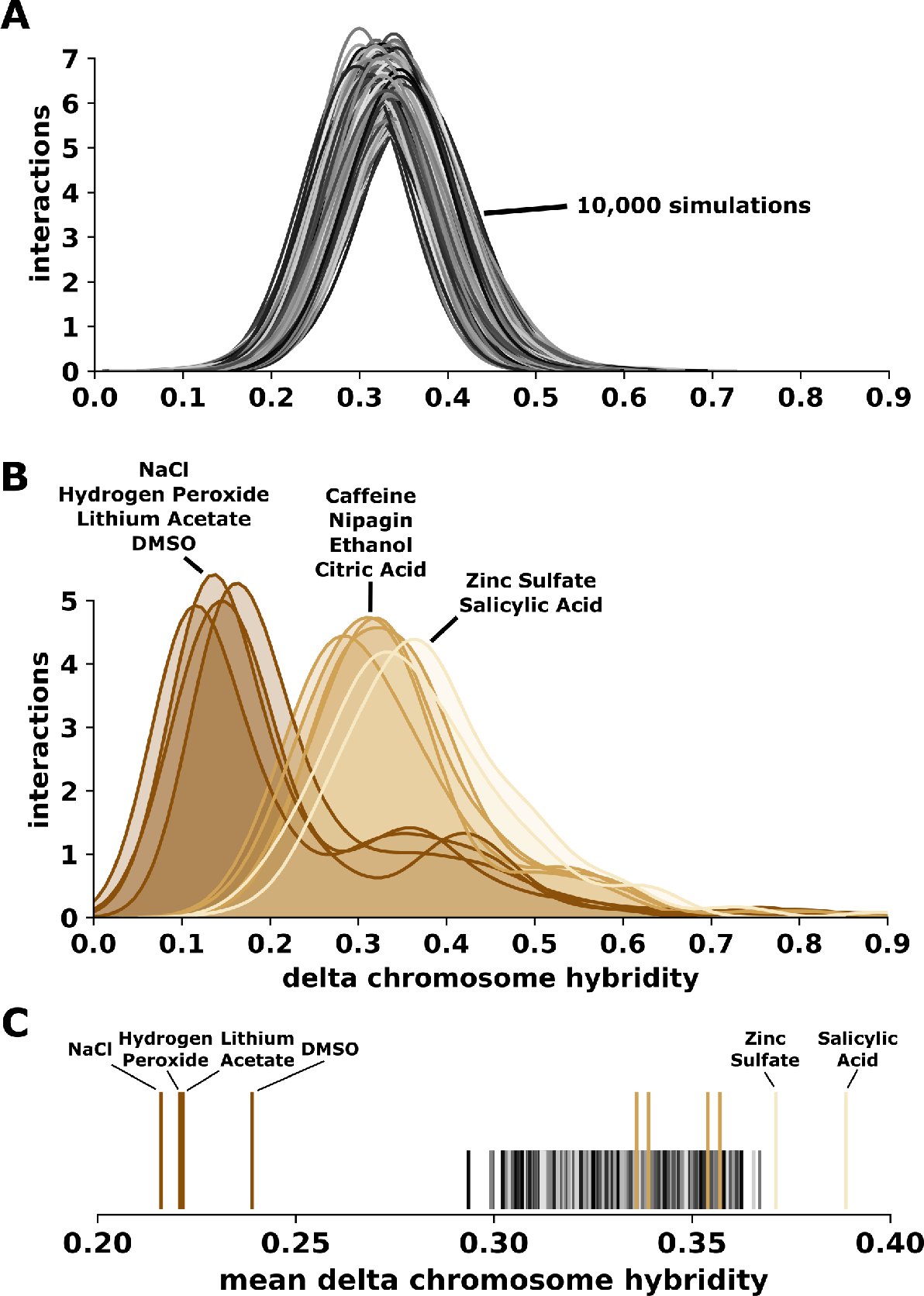
Delta chromosome hybridity (DCH) of simulated chromosomes. **(A)** The distribution of DCH in a simulated environment resulting from simulated chromosomes based on no chromosomal interactions. Each environment consists of 24 genomes, each consisting of 16 simulated chromosomes. Each simulated environment was repeated 10,000 times. **(B)** The distribution of DCH found in our experimental environments. Distributions are colored according to their mean DCH. **(C)** Mean DCH of our experimental environments as compared to the range of mean DCH found in simulations (grey-black lines).

### Genotype x Environment Interactions and Gene Ontology

Sampling all F2 hybrid genomes, a total of 315 10kb bins fixed for either *S. cerevisiae* or *S. paradoxus* were detected (406 bins including chromosome 5 and 7; Figure S6A). We found large variation between environments ranging from 36 regions in salicylic acid to 372 regions in NaCl. Seventy-eight of these regions were environment-specific, i.e. they were inherited from either *S. cerevisiae* or *S. paradoxus* across all 24 hybrid genomes per environment, but only found in one environment.

We identified 1884 genes that were either located in or overlapped with the 315 fixed regions, ranging from 51 genes in salicylic acid to 1533 genes in NaCl (Figure S6C). The large differences between environments in the number of fixed alleles suggest that some stress conditions require more complex, quantitative adaptations with a polygenic basis, which is well-known to be the case for yeast adapting to salt for instance (Cubillos, et al. 2011; Dhar, et al. 2011).

Gene annotation analysis showed that most of the 1884 genes are involved in transport and pathways related to fungal cell wall organization (Figure S6B). Across environments, we found almost three times more genes fixed for the *S. cerevisiae* allele (n = 1373) than fixed for the *S. paradoxus* allele *(n* = 511). Only the F2 hybrid genomes isolated from salicylic acid showed more genes fixed for *S. paradoxus* (46 of 51 genes in total).

Only one region was consistently inherited from *S. cerevisiae* across all environments (chromosome 4: 970 000 - 980 000 bp, harbouring 5 genes), and one from *S. paradoxus* across all environments (chromosome 15: 10 000 - 20 000 bp, harbouring 7 genes). Among these are two general stress response genes, *HSP78* and *PAU20*. *HSP78* on chromosome 4 is associated with heat stress and mitochondrial genome maintenance (Leonhardt, et al. 1993; von Janowsky, et al. 2006). *PAU20* on chromosome 15 is associated with proteome stress and is upregulated during wine fermentation (Rossignol, et al. 2003; Marks, et al. 2008; Luo and van Vuuren 2009). It is surprising that the *S. paradoxus* allele of *PAU20* became fixed across all environments as we normally expect *S. cerevisiae* to be better adapted to fermentation environments.

We did not identify fixations of alternative alleles at the same locus in different environments (e.g. a bin that was consistently inherited from *S. cerevisiae* across all 24 hybrid genomes in ethanol but from *S. paradoxus* in salicylic acid).

**Figure 5.**
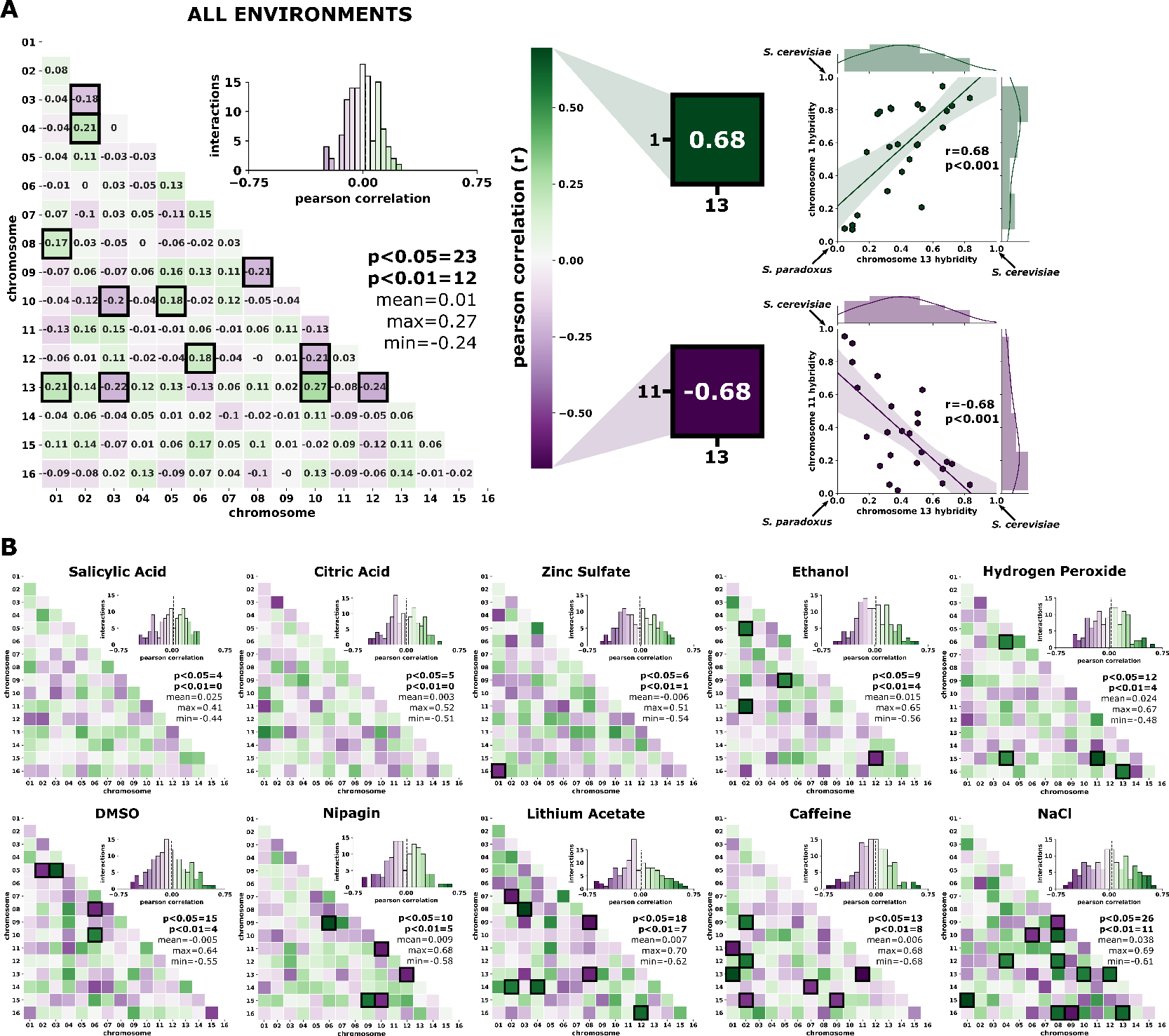
Epistatic interactions between chromosome hybridity across environments. **(A)** Pearson correlation coefficient (r) of chromosome hybridity (percent chromosome mapping to *S. cerevisiae*) between chromosomes across all environments (n genomes = 237). Inset depicts the distribution of Pearson correlation coefficients for all chromosome interactions. Positive correlations are shown in green and negative correlations are shown in purple. Statistically significant correlations (p < 0.01) are highlighted in black. Examples of strong correlations are shown for the interactions between chromosome 13 and 11 (positive) and chromosome 13 and 1 (negative) in the Caffeine environment. **(B)** Pearson correlation between chromosome hybridity within each environment. Calculations are performed as in panel B but for each environment independently (n ~ 24 genomes). Environments are sorted from top left to bottom right according to the number of significant (p < 0.01) correlations found.

## Discussion

Hybrid fitness varies among genotypes, generations, and environments (e.g. Nolte and Tautz 2010; Arnold, et al. 2012; Stelkens, Brockhurst, Hurst, Miller, et al. 2014; Hwang, et al. 2016b) but the genetic basis of hybrid fitness is usually unknown, especially in non-domesticated organisms (but see Rieseberg, et al. 1999; Payseur and Rieseberg 2016). In addition, the interaction between the environment and the targets of selection in hybrid genomes remains almost entirely unexplored (but see Shahid, et al. 2008; Hwang, et al. 2016a). Whether or not the variation contained in hybrid populations contributes to adaptive evolution crucially depends on the type of genetic mechanism underlying fitness. Under heterozygote advantage, for instance, the most fit genotype cannot breed true in sexual, diploid populations because segregation will inevitably break up beneficial allelic combinations (Buerkle and Rieseberg 2008). But if the most fit genotype is homozygous for alleles derived from each lineage at different loci, such recombinant homozygotes can breed true, may become fixed through drift or selection, and potentially establish new lineages (Fitzpatrick and Shaffer 2007).

Here, we describe the composition of 237 stress-resistant F2 hybrid genomes, made from two divergent *Saccharomyces* yeast species. We measured variance in overall hybridity, interspecific heterozygosity, and epistasis found between and within hybrid genomes. We found that the composition of hybrid genomes was strongly contingent on environmental context. Genomes from different environments varied in every aspect of hybridness measured. First, genomes from different environments differed in the proportion of interspecific heterozygosity. While some environments clearly selected for more homozygous genomes (e.g. zinc sulfate), others selected for more heterozygous genomes (e.g. NaCl; Figure 1B). Second, individual chromosomes exhibited strong species biases depending on the type of stress they were exposed to (Figure 2B). This is presumably because genes important for resistance to the specific toxin are located on these chromosomes, and alleles from parental species differ in how well they tolerate this toxin (but note that we did not observe fixation of opposite species alleles in different environments; Figure S6). Third, we found evidence for environment-dependent, non-homologous chromosomal associations (Figure 3). Positive same-species and to our surprise, also positive opposite-species chromosome associations occurred, depending in frequency and type on the environment (Figure 5). Computer simulations in a selection- and epistasis-free space with independently segregating chromosomes (Figure 4, S3, S4, S5) produced a smaller number of significant associations between chromosomes in hybrid genomes, suggesting that some heterospecific chromosomal combinations in experimental hybrid genomes were indeed under positive epistatic selection.

The presence of beneficial interspecific interactions between chromosomes from species with such large evolutionary divergence (~15%) is surprising. At this genetic distance, which is roughly three times larger than the distance between humans and chimpanzees, one would expect hybrids to be ridden with Dobzhansky-Muller incompatibilities (negative epistasis) and other negative fitness effects of species divergence, such as the missegregation of chromosomes due to antirecombination (Hunter, et al. 1996; Greig, et al. 2003; Rogers, et al. 2018). By virtue of our experimental design we only sampled from an already viable subset of F2 hybrids. Still, even in this subset this we would expect selection to favour same-species chromosomal combinations (in the resolution possible given limited rounds of segregation and recombination in the F2 generation).

Interestingly, when screening for recessive incompatibilities in the same hybrid cross, a previous study found that replacing chromosomes 4, 13, and 14 in *S. cerevisiae* with homologous chromosomes from *S. paradoxus* did not yield any viable haploids (Greig 2007). In our experiment, focusing here on the NaCl environment because it revealed the most significant epistatic interactions, the same three chromosomes (4, 13, 14) were involved in five of eight positive same-species interactions, suggesting that different-species combinations with any of these three chromosomes may indeed cause low fitness in hybrid genomes, and are selected against (Figure 5B). In addition, we found the three positive opposite-species associations in NaCl involving five chromosomes (6, 8, 9, 10, 16). Of these, the previous study found four chromosomes (6, 8, 9, 10) not to be problematic to transfer into a different species background (there was no suitable auxotrophic marker for chromosome 16, so this was a technical limitation, not an incompatibility), suggesting positive heterospecific epistasis, or at least an absence of incompatibilities on these chromosomes.

Individual chromosomes also differed in in their level of interspecific heterozygosity independently of the environment. For instance, chromosomes 4, 6, and 14 were more beneficial to fitness as homozygotes, while chromosomes 2, 3 and 12 were more beneficial as heterozygotes (Figure 1C). Overall however, interspecific heterozygosity was unexpectedly low across all hybrid genomes (Figure S1), suggesting that same-species allelic combinations (homozygosity) in an otherwise hybrid genomic background can provide high fitness. These results speak against strong effects of dominance or overdominance to hybrid fitness and are more consistent with selection against heterozygotes (underdominance), which has been shown in *Saccharomyces* (Laiba, et al. 2016), natural *Populus* (Lindtke, et al. 2012) and *Helianthus* hybrid populations (Lai, et al. 2005). Yet, dominance complementation of recessive alleles and overdominant interactions within loci are often reported from laboratory crosses of *Saccharomyces* where heterozygotes are fitter than both homozygotes due to simple or reciprocal complementation of one or several recessive deleterious mutations (Zörgö, et al. 2012; Plech, et al. 2014; Shapira, et al. 2014; Blein-Nicolas, et al. 2015; Laiba, et al. 2016). In some cases, this inconsistency with our results may be explained by the much smaller genomic divergence (usually <1%) between parental yeast lineages used in these studies (mostly all *S. cerevisiae* strains). In agreement with our results, a study on heterosis in the same interspecific yeast cross (*S. cerevisiae x S. paradoxus)* suggested that heterosis in hybrids was likely a result of both dominance complementation of recessive deleterious alleles, and additional overdominant or epistatic effects.

The excess of homozygosity in our experiment is consistent with many if not most high fitness alleles being recessive and only becoming expressed in the homozygous state. Alternatively, selection on some aspects of fitness in the haploid phase (during sporulation and germination) may have caused strains with the same beneficial combinations of parental chromosomes to mate, resulting in homozygous diploids. For instance, strains may be able to maximize their fitness by germinating early (Miller and Greig 2014), which could result in a process similar to assortative mating.

We acknowledge that the large size of the populations we used here, the facultative asexual reproduction and the ability of yeast to self-fertilize (hybrids do not rely on backcrossing), can help restore fitness quickly after outbreeding depression in the F2 hybrid generation. Together, this can catalyze the propagation of hybrid genotypes even with small and temporary fitness advantages, which is not a likely outcome in small populations of purely sexual species. It is also possible that some of the hybrids in our experiment were aneuploid, i.e. they lost or gained one or more chromosomes. Aneuploidy is a common by-product of chromosomal mis-segregation in the F1 hybrid meiosis of interspecific yeast crosses (Hunter, et al. 1996), and gene dosage effects ensuing from higher copy numbers may have helped some hybrids in our experiment survive stress. Aneuploidy has been associated with stress resistance in yeast and other fungi (Selmecki, et al. 2009; Pavelka, et al. 2010; Kwon-Chung and Chang 2012), and has been suggested to serve as transient adaptation mechanism (Chang, et al. 2013; Hose, et al. 2015; Smukowski Heil, et al. 2016).

Hybridization mostly occurs at the margins of species ranges where conditions are more extreme than in the center of their distribution, i.e. in more stressful environments. Hybridization also occurs more frequently in human- or otherwise perturbed habitats, where geographic and ecological species barriers are lost (King, et al. 2015; Arnold 2016; Gompert and Buerkle 2016; McFarlane and Pemberton 2019). Thus, the circumstances leading to hybridization often coincide with times of increased environmental stress, which in turn creates ecological opportunity. The large number of significant chromosome-by-environment interactions we found in our hybrid populations showcases the genetic variation characteristic of hybrid swarms. This can facilitate their functional diversification and, potentially, the colonization of novel and challenging environments as shown for instance in *Helianthus* sunflowers and African cichlid fish (Rieseberg, et al. 2003; Seehausen 2004; Nolte, et al. 2005; Abbott, et al. 2013). But in a genome that is a patchwork of heterozygous and homozygous regions, knowing the type of genetic mechanism responsible for increased fitness is essential for any predictions regarding the role of hybridization in adaptive evolution. In addition, our data suggests that each environment selects for different parental alleles and for different epistatic interactions.

In conclusion, both environmental and genetic contingencies of hybrid fitness limit our ability to predict the evolutionary and ecological outcomes of hybridization (Gompert and Buerkle 2016). For instance, hybrids could outcompete and drive a parent population to extinction in one locality, but the same hybrids may have inferior fitness in a different habitat, posing no threat at all to the parents. The results of our study show that in order to capture the risks and benefits of genetic exchange between populations and species, it is important to measure multiple dimensions of hybridity and measure hybrid fitness in multiple environmental contexts.

## Materials and Methods

### Parent and F1 hybrid strains

We chose two genetically tractable laboratory strains as parents: *S. cerevisiae* haploid strain YDP907 (*MATα, ura3::KanMX, can1r*), isogenic with strain background S288c, was crossed to *S. paradoxus* strain YDP728 (*MATa, ura3::KanMX, cyh2r*), which is isogenic with strain background N17. This produced a diploid F1 hybrid (MATa/@, *ura3::KanMX, cyh2r, can1r*), which was purified by streaking and then stored frozen at −80°C in 20% glycerol stock.

### F2 hybrids

To make F2 hybrids, we grew a 5ml culture of the F1 hybrid in YEPD (1% yeast extract, 2% peptone, 2% dextrose) for 24h at 30°C, and then transferred 200µl of this to 50ml of KAC sporulation medium (1% potassium acetate, 0.1% yeast extract, 0.05% glucose, 2% agar) which was shaken at room temperature for 5 days to induce meiosis and sporulation. Sporulation was confirmed under the microscope. 50µl of the spore culture, in serial dilutions and using three replicates per dilution, were then plated onto arginine drop out agar plates supplemented with the drugs canavanine (60mg/L) and cycloheximide (3.33mg/L). These drugs kill the fully heterozygous F1 hybrids because the resistance alleles *cyh2r* and *can1r* are recessive. But those meiotic spores (produced by the F1 hybrid) that contain both resistance alleles can, if viable, germinate and form colonies. This method ensured that only F2 hybrids were sampled and sequenced.

### Environments and stress

50µl of spore culture (containing 25.5 double drug-resistant viable spores on average confirmed by streaking out on double-drug medium) were used as the founding population for inoculation and growth on flat-bottomed, 96-well cell culture plates. Wells contained liquid minimal medium plus uracil, cycloheximide and canavanine (0.67% yeast nitrogen base without amino acids, 2% glucose, 2% agar, 0.003% uracil, 60 mg/L canavanine, 3.33mg/L cycloheximide), allowing for the germination, mating, and growth of the double drug-resistant F2 progeny. This was supplemented with a range of concentrations of ten toxins (one at a time): salicylic acid (C_7_H_6_O_3_), caffeine (C_8_H_10_N_4_O_2_), ethanol (C_2_H_6_O), zinc sulfate (ZnSO_4_), hydrogen peroxide (H_2_O_2_), methyl paraben (“Nipagin”; CH_3_(C_6_H_4_(OH)COO), sodium chloride (NaCl), lithium acetate (CH_3_COOLi), dimethyl sulfoxide (“DMSO”; C_2_H_6_OS), and citric acid (C_6_H_8_O_7_). Toxicity gradients (or ‘environmental clines’) were generated along the y-axis of the 96-well plates such that the bottom row contained the lowest concentration, and the top row contained the highest concentration of the toxin, lethal for all strains (exact concentrations in Table S1).

Plates were incubated at 30°C for 4 days. Then, 1µl from each well was transferred to the same position on a new 96-well culture plate containing identical concentrations. The optical density (OD) of each well was measured with a microplate reader (Infinite M200 Pro, Tecan) (time point t_0_) and plates were incubated at 30°C for another 3 days. After this, another OD measurement was taken (t_1_) and plates were stored at 4°C until further processing. We calculated, for each well, whether growth had occurred by subtracting ODt_0_ from ODt_1_. Assuming that a doubling in optical density approximately equals a doubling of cell numbers, cells in permissive environments at the bottom of the plate completed on average 3 cell cycles (2.78 ± 0.15 across toxins) from t_0_ to t_1_ whereas cells in the topmost row did not divide at all.

We sampled from all twelve wells of the bottom row (i.e. the lowest stress) and from the twelve wells of the highest stress that allowed growth from each column, from all ten plates. These 240 samples were streaked out for single colonies on YEPD plates and grown for 48h at 30°C. We then picked a single colony from each sample, and froze it for sequencing.

### DNA extraction, PCR and sequencing protocols

DNA was extracted using the MasterPure Yeast DNA Purification Kit (Epicentre). A genotyping-by-sequencing protocol was modified from microsatellite library preparation and ddRAD sequencing approaches as follows (Nolte, et al. 2005; Peterson, et al. 2012). The library construction is based on an efficient combined restriction digest/adaptor ligation. The restriction enzymes Csp6I (which cleaves 5’- G^T A C -3’ sites) and PstI (which cleaves 5’- C T G C A^G -3’ sites) were used to digest genomic DNA to generate sticky ends. The reaction conditions permit that sticky end adapters and T4 ligase are added to the reaction such that adaptors are ligated to the restriction sites. Importantly, the adaptors do not fully reconstitute the restriction sites. Thus, once an adaptor is ligated, this site will not be recut while an undesired ligation of two genomic DNA sticky ends will be recut until all DNA ends are saturated with an adaptor. For this purpose we modified the ligation sites of the Illumina Truseq adapters such that they matched the sticky ends generated by the restriction enzymes. Further, we labeled the adapters to include 24 and 16 different molecular barcodes (MIDs) respectively, which could be combined in 384 different combinations for multiplexed sequencing (paired ends) of the libraries on an Illumina Myseq sequencer.

### Quality filtering of raw reads and mapping protocol

We examined the quality of raw ddRADseq reads of each sample using FastQC (v0.11.8) (FastQC 2018). Illumina sequencing adapters, primer sequences, ambiguous and low quality nucleotides (PHRED quality score < 20) were removed from both paired-end reads according to the default parameters in the NGS QC Toolkit (v2.3.3) (Patel and Jain 2012). Three of 240 F2 genomes were abandoned due low sequence yield. A total of 8.2 Gb high-quality reads were generated after quality control with mean depth of 12.84 per marker.

Parental strain sequences were downloaded from the *Saccharomyces* Genome Database website. The S288c reference genome used was R64-2-1_20150113. The N17 reference genome was obtained from ftp://ftp.sanger.ac.uk/pub/users/dmc/yeast/latest/, in the para_assemblies/ N_17 folder.

Given the high sequence divergence between *S. cerevisiae* and *S. paradoxus* (15%), correct assignment of reads to the parental species was efficient using NovoAlign (NovoAlign 2018) with default parameters. For this, both parental genomes were concatenated and used as a combined reference for mapping, and species affiliations and chromosomal positions of successfully mapped reads were written to the same output file. Mapped reads were considered correctly assigned to the reference sequence when the mapping quality was >= 20, indicating a single and unique best match.

### Ploidy determination

To differentiate haploid and diploid F2 hybrid genomes (to be able to score the prevalence of homo- and heterozygosity), we mapped paired-end reads again, this time only to the *S. cerevisiae* S288c reference genome using NovoAlign (NovoAlign 2018) with the default parameters. We sorted mapped files according to their genomic coordinates using Picard Tools v2.18.23 (Picard 2019) and performed variant calling in FreeBayes (Garrison and Marth 2012) with at least five supporting reads required to consider a variant. SNPs were selected and filtered using GATK (Li, et al. 2008) according to stringent filtering criteria with the following settings: a) QUAL >30.0; b) QD >5.0; c) FS <60.0; d) MQ >40.0; e) MQRankSum >-12.5; and f) ReadPosRankSum >- 8.0. Additionally, if there were more than 3 SNPs clustered in a 10-bp window, we considered all three SNPs false positives and removed them. After filtering, we identified 14,975 SNPs, ranging between 10,725 and 12,345 among environments.

Haploidy or diploidy was called using the heterozygosity score of biallelic SNPs using a custom bash script. Theoretically, both the number of heterozygous SNPs and ratio of heterozygous SNPs over the total SNPs in haploids should be zero (except in case of sequencing errors and aneuploidies). However, the absolute number of heterozygous SNPs is affected by the overall sequence size of each sample, and the ratio is affected by the number of SNPs derived from *S. paradoxus* (because of the large nucleotide divergence between the parents, ~15% of the reads from *S. paradoxus* do not map to the *S. cerevisiae* reference genome). Also, although unlikely, diploid F2 genomes can theoretically be fully homozygous with both copies of eight chromosomes inherited from each parental species, assuming genetic material from both parental species is equally represented in the F2 generation. At zero heterozygosity, haploid and diploid genomes are indistinguishable by SNP calling and mapping to parental genomes. Thus, at the risk of omitting extremely homozygous diploids, we applied two criteria to consider a genome diploid: 1) The number of heterozygous SNPs in a diploid F2 genome should be larger than the average number of heterozygous SNPs found in both parental genomes. 2) The ratio of heterozygous over hetero-plus homozygous SNPs in a diploid F2 genome should be larger than the average ratio of heterozygous over hetero-plus homozygous SNPs in parental genomes. Since there are nearly zero heterozygous SNPs in the *S. cerevisiae* genome, this can be simplified to:

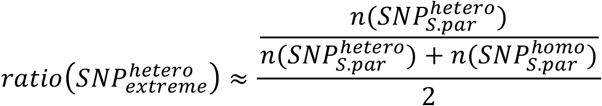

Using this analysis, we assigned 187 of the 237 F2 genomes to be diploid.

### Chromosome zygosity

To determine the zygosity of each chromosome in diploid F2 genomes, we mapped high quality reads of each sample to the reference combining both parental genomes. We extracted the species affiliation of each read using a custom bash script. The chromosomal content per species per sample was extracted using custom perl code by calculating the proportion of markers mapping to only one species over markers mapping to both species. We considered chromosomes heterozygous if marker proportions fell between 0.25 – 0.75. Chromosomes with smaller or larger species content were assigned homozygous.

To understand what causes variation in zygosity between environments, chromosomes and genomes, we modeled chromosome heterozygosity using a mixed-effect linear model. Chromosome zygosity was used as the response variable in the model and was predicted as a function of fixed predictors environment and chromosome ID and their interactions and sample ID as a random effect. We selected the most appropriate model by identifying the simplest model that maintained the lowest Akaike Information Criteron (AIC; (Akaike 1974)). AIC optimizes the relationship between the fit and complexity of a model by balancing the fit of the model with the number of parameters estimated (Harrison, et al. 2018).

### Genome-level and chromosome-level hybridity

To describe the genome-level and chromosome-level hybridity we calculated the percentage of the entire genome or chromosome for each sample that mapped to either the *S. cerevisiae* or the *S. paradoxus* genome. We defined hybridity as the percent of markers per genome or chromosome mapping to *S. cerevisiae (*100% = *S. cerevisiae*, 0% = *S. paradoxus*). This was calculated across all environments, as well as for each environment individually. We then used mixed-effect linear models with genome and chromosome hybridity as response variables. We predicted hybridity to be a function of the fixed effects *environment, chromosome ID, ploidy* and their interactions, and included sample ID as random effect. We selected the most appropriate model by starting with the full model and removing insignificant components, identifying the simplest model that maintained the lowest AIC.

To characterize the interactions between chromosomes within a F2 genome, we calculated the difference in chromosome hybridity for each chromosome combination, defined as delta chromosome hybridity (DCH). Large DCH indicates that the two chromosomes map primarily to opposing species within a genome. Small DCH indicates that the chromosomes have similar levels of hybridity, suggesting that either the chromosomes come from primarily the same species or at least have similar hybridity proportions. DCH was calculated for all environments together, as well as each environment individually. To investigate the epistatic interactions between chromosomes we determined the Pearson correlation (r) between chromosome hybridity for all pairwise chromosome combinations. A significant positive correlation between two chromosomes suggests a positive epistatic interaction within a species, meaning that as chromosome hybridity in one chromosome shifts towards one species (1 = *S. cerevisiae*, 0 = *S. paradoxus*), the chromosome hybridity in the linked chromosome also shifts towards the same species. A significant negative correlation between two chromosomes suggests a negative epistatic interaction within a species. In this case, as one chromosome shifts towards one species, the linked chromosome shifts towards the other species.

### Simulated chromosomes

In order to determine if the observed zygosity, hybridity and chromosomal interactions found in each environment deviated from neutral expectations, we developed simulations that determined the expected range for these measurements based solely on chance, assuming no selection and free segregation of chromosomes in hybrid genomes. Each simulation displayed the expected distribution of zygosity, hybridity or chromosomal interactions that could be found in a selection-free environment with no epistasis between chromosomes. Each simulated environment was repeated 10,000 times. Each environment consisted of 24 genomes and each genome consisted of 16 chromosomes as found in our data. Each chromosome was randomly assigned a chromosome hybridity score between 0 and 1. To study the expected zygosity, chromosomes were labeled as either homozygous for *S. cerevisiae* or *S. paradoxus* or heterozygous based on the same rules used in the analysis of our data (see ****Chromosome zygosity****). The heterozygosity and hybridity for each genome were then calculated for each simulation. The means and distributions of these values were plotted and compared to the means and distributions we received from our data (Figure S1, S3). To study the expected chromosome interactions, we calculated the 120 interactions that occur within a simulated genome between chromosomes. Delta chromosome hybridity (DCH) and Pearson correlation (r) were calculated in the same manner as our data was analyzed (see ****Genome-level and chromosome-level hybridity****). The distributions of these interactions were then plotted and compared to the data from our study (Figure 4 and S4). For the Pearson correlation (r), we also determined the range of expected significant positive and negative correlations and compared them to our data (Figure S5).

### Genotype x Environment Interactions and Gene Ontology

To explore for gene-by-environment interactions, i.e. which gene from which species background is more frequently found in which environment, we divided genomes into 10kb bins. Assuming each bin is a locus and the parental species affiliations are the two possible alleles, we scored bins as 1 if all markers within that bin mapped to the *S. cerevisiae* reference genome, and 0 if all markers mapped to the *S. paradoxus* reference genome across all 24 genomes within a given environment (excluding chromosome 5 and 7). Genes located in or overlapping with these fixed regions of each environment were further inspected using the gene annotation data base DAVID (Huang, et al. 2009). Environment-independent fixations of alleles (bins fixed for the same species across all environments) and unique fixations (bin fixed only for one environment) were extracted using custom R code.

## Author contributions

RS and DG received the project idea and designed the experiment. RS performed the experiment and collected the samples. AWN planned and performed the library preparation and sequencing. ZZ processed the ddRADseq data. ZZ, DPB and TJ analyzed the data. RS, ZZ and DPB wrote the manuscript.

## Acknowledgements

We thank Elke Bustorf for the preparation of libraries and David Rogers for advice on mapping. This work was supported by the Max Planck Society (to DG), ERC starting grant (“EVOLMAPPING” to AWN), Swedish Vetenskapsrådet (grant number 2017-04963 to RS), Carl Trygger foundation (to ZZ) and Wenner-Gren Foundations (grant number UPD2018-0196 to DPB).

## Supplementary Material

**Table S1.** Toxins and concentrations. Concentrations (in M or %) of the 10 toxins used in the experiment. Numbers 1-8 indicate row numbers from top (lethal) to bottom (permissive) on 96-well culture plates.

| Substance |  | Concentration |  |  |  |  |  |  |  |
| --- | --- | --- | --- | --- | --- | --- | --- | --- | --- |
|  |  | 1 | 2 | 3 | 4 | 5 | 6 | 7 | 8 |
| salicylic acid (C<sub>7</sub>H<sub>6</sub>O<sub>3</sub>) | M | 0.086 | 0.057 | 0.038 | 0.025 | 0.017 | 0.011 | 0.008 | 0.005 |
| caffeine (C<sub>8</sub>H<sub>10</sub>N<sub>4</sub>O<sub>2</sub>) | M | 0.103 | 0.069 | 0.046 | 0.031 | 0.020 | 0.014 | 0.009 | 0.006 |
| zinc sulfate (ZnSO<sub>4</sub>), | M | 0.186 | 0.124 | 0.083 | 0.055 | 0.037 | 0.024 | 0.016 | 0.011 |
| Nipagin (CH<sub>3</sub>(C<sub>6</sub>H<sub>4</sub>(OH)COO)) | M | 0.053 | 0.035 | 0.024 | 0.016 | 0.010 | 0.007 | 0.005 | 0.003 |
| sodium chloride (NaCl) | M | 4.0 | 2.667 | 1.778 | 1.185 | 0.790 | 0.527 | 0.351 | 0.234 |
| citric acid (C<sub>6</sub>H<sub>8</sub>O<sub>7</sub>) | M | 0.115 | 0.077 | 0.051 | 0.034 | 0.023 | 0.015 | 0.010 | 0.007 |
| lithium acetate (CH<sub>3</sub>COOLi) | M | 0.5 | 0.333 | 0.222 | 0.148 | 0.099 | 0.066 | 0.044 | 0.029 |
| hydrogen peroxide (H<sub>2</sub>O<sub>2</sub>) | % | 0.014 | 0.009 | 0.006 | 0.004 | 0.003 | 0.002 | 0.001 | 0.001 |
| ethanol (C<sub>2</sub>H<sub>6</sub>O) | % | 50 | 33.333 | 22.222 | 14.815 | 9.877 | 6.584 | 4.390 | 2.926 |
| DMSO (C<sub>2</sub>H<sub>6</sub>OS) | % | 30 | 20.000 | 13.333 | 8.889 | 5.926 | 3.951 | 2.634 | 1.756 |

**Table S2.**
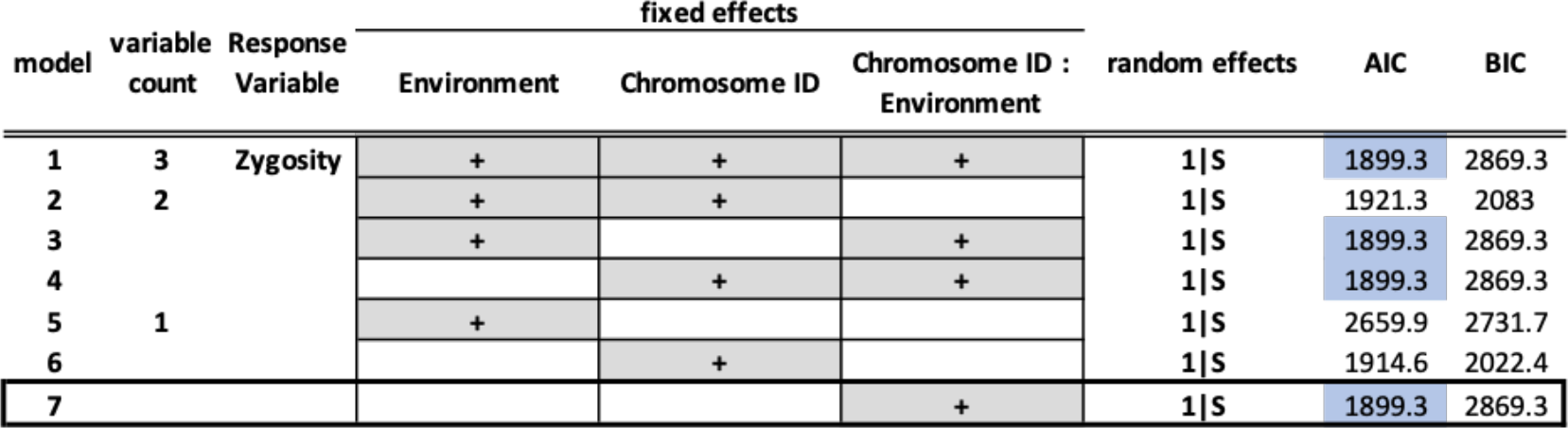
Linear mixed-effect models used to predict zygosity. The most appropriate model was selected by identifying the simplest model that maintained the lowest Akaike Information Criteron (AIC). The optimal model, given our data, is indicated by a black square.

**Table S3.** Linear mixed-effect models used to predict chromosome hybridity. The most appropriate model was selected by identifying the simplest model that maintained the lowest Akaike Information Criteron (AIC). The optimal model, given our data, is indicated by a black square.

| model | variable count | Response Variable | fixed effects |  |  |  |  | random effects | AIC | BIC |
| --- | --- | --- | --- | --- | --- | --- | --- | --- | --- | --- |
|  |  |  | Environment | Chromosome ID | Ploidy | Chromosome ID : Environment | Chromosome ID : Ploidy |  |  |  |
| 1 | 5 | Hybridity | + | + | + | + | + | 1\|S | -742.4 | 368.4 |
| 2 | 4 |  | + | + | + | + |  | 1\|S | -742.4 | 368.4 |
| 3 |  |  | + | + | + |  | + | 1\|S | -672.1 | -403.8 |
| 4 |  |  | + | + |  | + | + | 1\|S | -742.4 | 368.4 |
| 5 |  |  | + |  | + | + | + | 1\|S | -742.4 | 368.4 |
| 6 |  |  |  | + | + | + | + | 1\|S | -742.4 | 368.4 |
| 7 | 3 |  | + | + | + |  |  | 1\|S | -644.8 | 470 |
| 8 |  |  | + | + |  | + |  | 1\|S | -727.2 | 283.8 |
| 9 |  |  | + | + |  |  | + | 1\|S | -672.1 | -403.8 |
| 10 |  |  | + |  | + | + |  | 1\|S | -726.1 | 291.2 |
| 11 |  |  | + |  | + |  | + | 1\|S | -672.1 | -403.8 |
| 12 |  |  | + |  |  | + | + | 1\|S | -742.4 | 368.4 |
| 13 |  |  |  | + | + | + |  | 1\|S | -726.1 | 291.2 |
| 14 |  |  |  | + | + |  | + | 1\|S | -664.8 | -452.6 |
| 15 |  |  |  | + |  | + | + | 1\|S | -742.4 | 368.4 |
| 16 |  |  |  |  | + | + | + | 1\|S | -742.4 | 368.4 |
| 17 | 2 |  | + | + |  |  |  | 1\|S | -645.9 | -477.4 |
| 18 |  |  | + |  | + |  |  | 1\|S | 1109.7 | 1190.8 |
| 19 |  |  | + |  |  | + |  | 1\|S | -727.2 | 283.8 |
| 20 |  |  | + |  |  |  | + | 1\|S | -672.1 | -403.8 |
| 21 |  |  |  | + | + |  |  | 1\|S | -637.4 | -518.4 |
| 22 |  |  |  | + |  | + |  | 1\|S | -727.2 | 283.8 |
| 23 |  |  |  | + |  |  | + | 1\|S | -664.8 | -452.6 |
| 24 |  |  |  |  | + | + |  | 1\|S | -726.1 | 291.2 |
| 25 |  |  |  |  | + |  | + | 1\|S | -664.8 | -452.8 |
| 26 |  |  |  |  |  | + | + | 1\|S | -742.4 | 368.4 |
| 27 | 1 |  | + |  |  |  |  | 1\|S | 1108.3 | 1183.2 |
| 28 |  |  |  | + |  |  |  | 1\|S | -634.8 | -522.5 |
| 29 |  |  |  |  | + |  |  | 1\|S | 1109.2 | 1134.1 |
| 30 |  |  |  |  |  | + |  | 1\|S | -727.2 | 283.8 |
| 31 |  |  |  |  |  |  | + | 1\|S | -664.8 | -452.6 |
| 32 | 0 |  |  |  |  |  |  | 1\|S | 1110.6 | 1129.3 |

**Figure S1.**
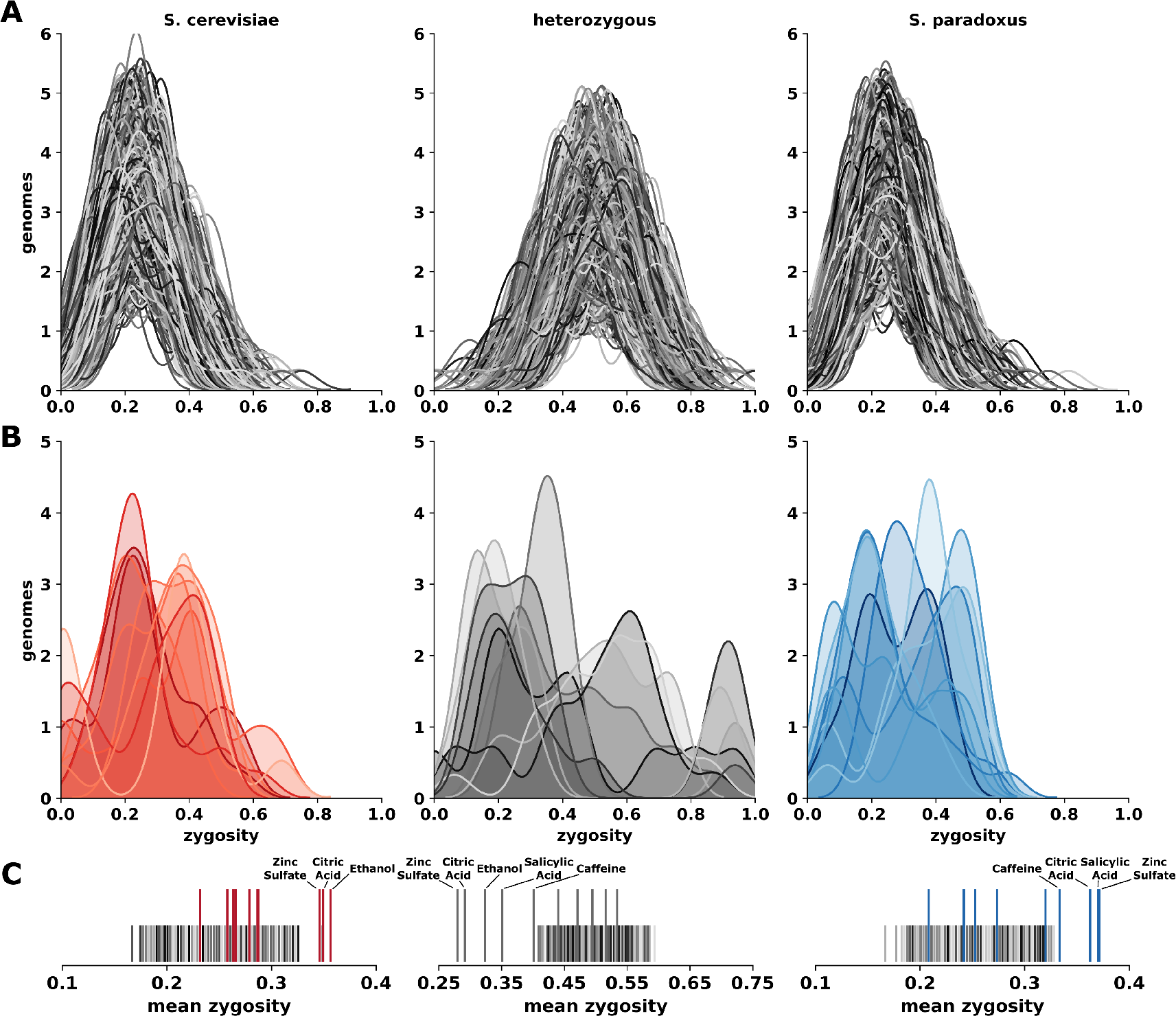
Genome-wide zygosity of simulated chromosomes. **(A)** The distributions of homozygosity for *S. cerevisiae* or *S. paradoxus* and heterozygosity in a simulated environment resulting from simulated chromosomes based on no chromosomal interactions (no epistasis) and random segregation. Each environment consists of 24 genomes, each consisting of 16 simulated chromosomes. Each simulated environment was repeated 10,000 times. **(B)** The distribution of zygosities found in our experimental environments. Distributions are colored according to their zygosity (red = homozygous *S. cerevisiae*, grey = heterozygous, blue = homozygous *S. paradoxus*). **(C)** Mean zygosity of our experimental environments as compared to the range of mean zygosities found in simulations (grey-black lines).

**Figure S2.**
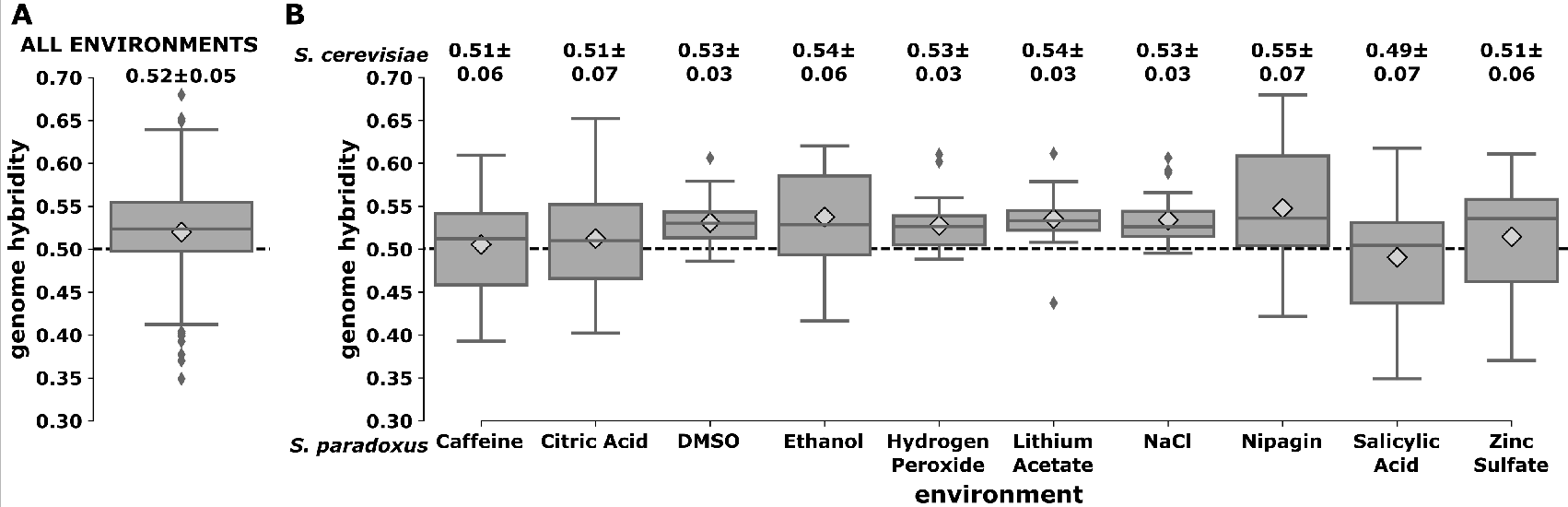
Genome-wide hybridity of 237 F2 hybrids. **(A)** Genome hybridity across all chromosomes and all environments, measured as the mean of the 16 chromosome hybridities (Figure 2) within a genome. Boxplots indicate the median and interquartile range. Diamonds indicate the mean. Dashed line (0.5) indicates equal amount of the genome mapping to *S. cerevisiae*. **(B)** Genome hybridity within environments. Numbers indicate the average genome hybridity for each F2 genome within each environment (n genomes ~ 24).

**Figure S3.**
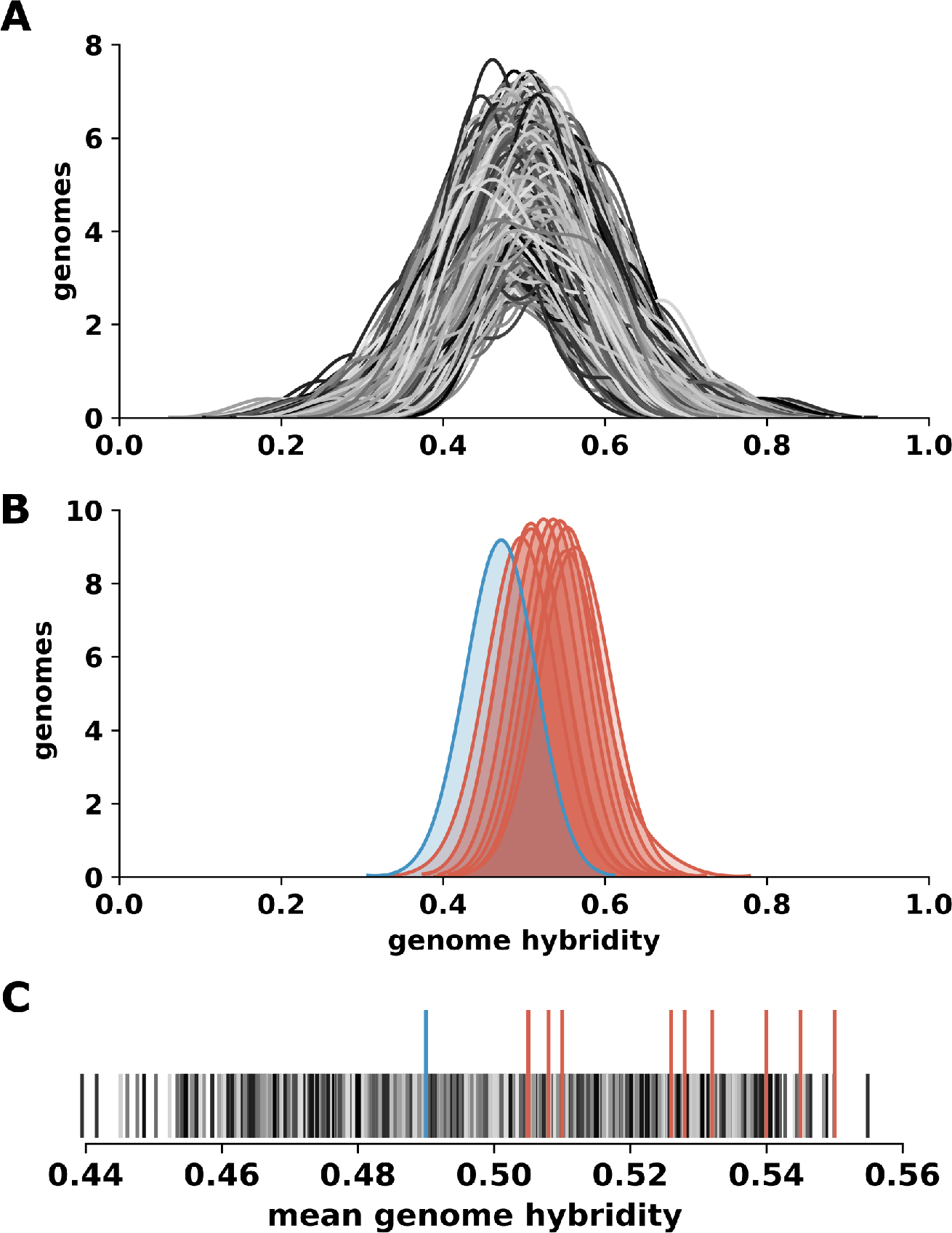
Genome-wide hybridity of simulated chromosomes. **(A)** The distribution of genome-wide hybridity in a simulated environment resulting from simulated chromosomes based on no chromosomal interactions (no epistasis). Each environment consists of 24 genomes, each consisting of 16 simulated chromosomes. Each simulated environment was repeated 10,000 times. **(B)** The distribution of genome-wide hybridity found in our experimental populations. Distributions are colored according to their overall mean species bias (red = *S. cerevisiae*, blue = *S. paradoxus*). **(C)** Mean genome-wide hybridity of our experimental populations as compared to the range of genome-wide hybridity found in simulations (grey-black lines).

**Figure S4.**
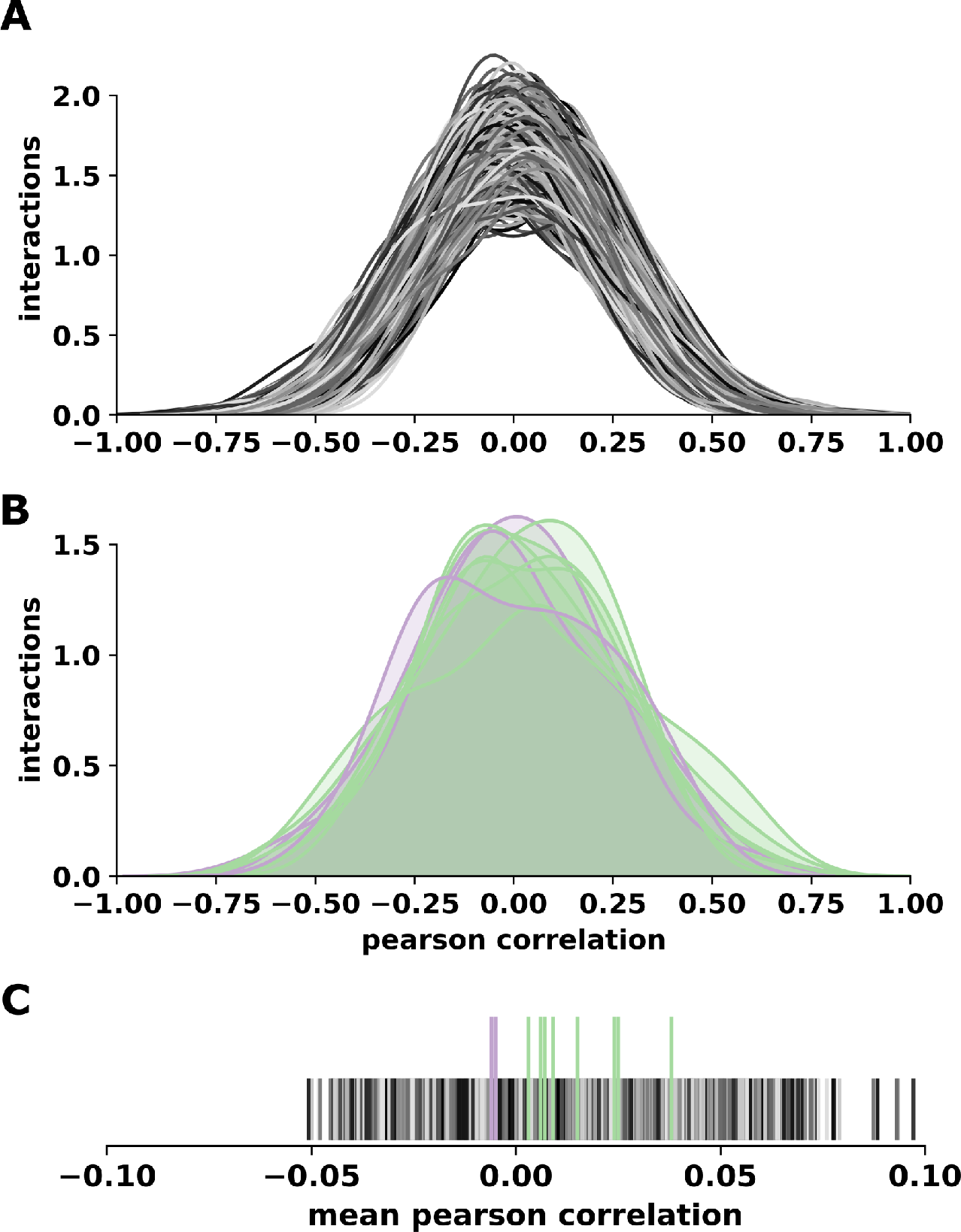
Pearson correlation (r) of simulated chromosomes. **(A)** The distribution of Pearson correlations in a simulated environment resulting from simulated chromosomes based on no chromosomal interactions (no epistasis). Each environment consists of 24 genomes, each consisting of 16 simulated chromosomes. Each simulated environment was repeated 10,000 times. **(B)** The distribution of Pearson correlations found in our experimental environments. Distributions are colored according to their mean Pearson correlation. **(C)** Mean Pearson correlation of our experimental environments as compared to the range of mean Pearson correlations found in simulations (grey-black lines).

**Figure S5.**
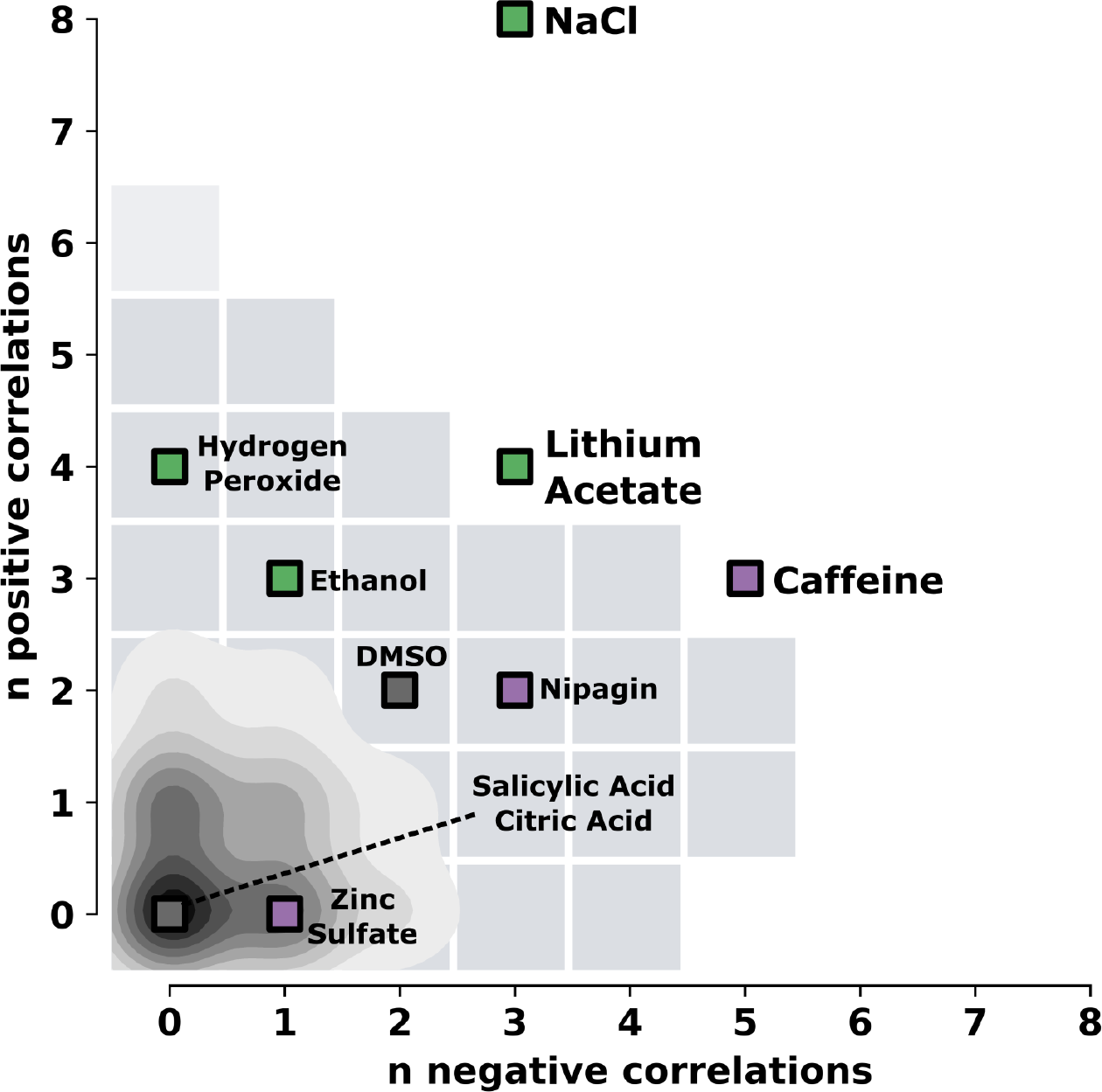
Distribution of significant (p < 0.01) positive and negative correlations resulting from simulated chromosomes. Each environment consists of 24 genomes, each consisting of 16 simulated chromosomes. Each simulated environment was repeated 10,000 times. A grey square indicates that at least one simulated environment resulted in that many negative and positive correlations. The density plot indicates the distribution of simulated environments. Our experimental environments are indicated with squares and labels. The color of each environment indicates the bias towards positive (green) or negative (purple) correlations.

**Figure S6.**
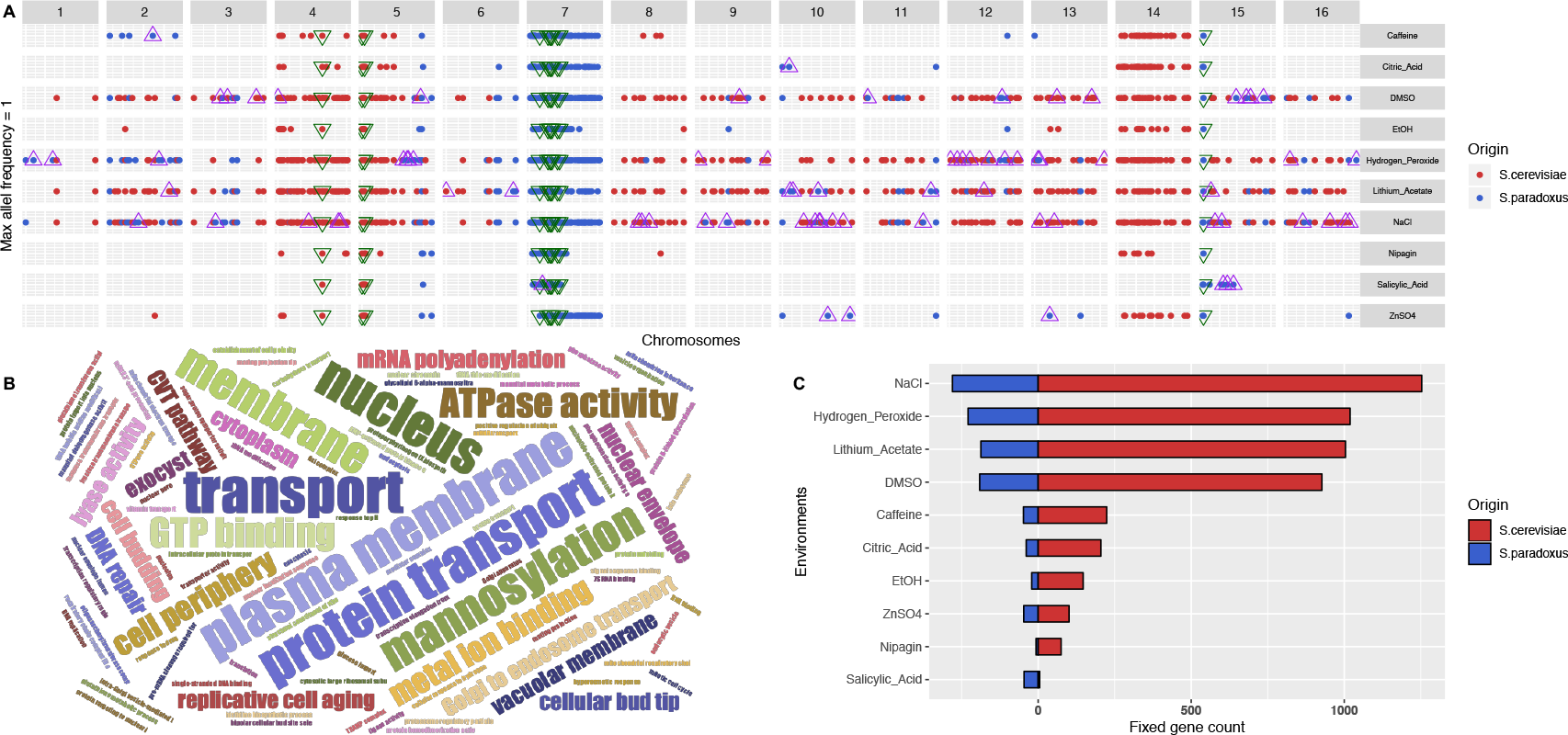
Genotype-by-environment interactions. **(A)** 10kb bins fixed for either one or the other parental allele across all genomes sampled from a given environment. All 16 chromosomes are shown. Bins fixed for the same species across all environments are marked with green triangles. Environment-specific regions fixed for the same species in all 24 genomes sampled from the same environment are marked with purple triangles. **(B)** Word cloud summarizing results of gene ontology analysis of the 1884 genes overlapping with or located within all fixed bins using DAVID (Huang, *et al.* 2009) **(C)** Bar chart showing the number of genes fixed for the *S. paradoxus* allele on the left and the *S. cerevisiae* allele on the right, by environment.

